# Gene expression and pollen performance indicate altered postmating selection between *Solanum* species with different mating systems

**DOI:** 10.1101/2024.10.31.621143

**Authors:** Timothy J Biewer-Heisler, Matthew JS Gibson, Emily Sornay, Leonie C Moyle

**Affiliations:** Department of Biology, Indiana University, Bloomington, Indiana USA, 1001 E 3rd St Room 325, Bloomington, IN 47405

**Keywords:** pollen-pistil interactions, gene expression, selection, cryptic female choice, postmating prezygotic, mating systems, pollen tube growth rate, *Solanum*

## Abstract

Postmating prezygotic (PMPZ) traits play an important role in mating success, especially in species where gametes from multiple males compete. Despite this, the effect of mating system transitions, and attendant shifts in the intensity of sexual selection, on specific PMPZ traits and their underlying loci is still poorly understood. Here we assessed differences in pollen PMPZ traits and tissue-specific gene expression (in leaf, pollen, and style) between two closely related plant species with different mating systems—*Solanum lycopersicum* (selfing) & *Solanum pennellii* (outcrossing). We focused on species differences in loci with known roles in pollen tube growth rate, including pectin methylesterases (PMEs) and their inhibitors (PMEIs), and *in-vitro* & *in-vivo* pollen tube growth rates. Among the gene expression differences observed between species, we found that the expression domain of pollen-biased genes was much narrower in the selfing species *S. lycopersicum* compared to the outcrossing species *S. pennellii*, including for most reproductive PMEs and PMEIs. In addition, *S. pennellii* had faster pollen tube growth rates *in-vivo*, while *S. lycopersicum* had faster *in-vitro* pollen tube growth rates. We propose that the lower expression of pollen tube development genes in *S. lycopersicum* style tissue, and reduced *in-vivo* pollen performance, is a result of reduced allocation to stylar mechanisms that modulate pollen tube growth, potentially consistent with relaxed selection on cryptic female choice in the selfing species.

**Article Summary:** This study investigates the evidence for differential postmating sexual selection in plants, by analyzing reproductive gene expression and pollen performance in two *Solanum* species with different mating systems. It finds that species differ systematically in pollen tube growth rates, and that pollen (‘male’)-biased genes have higher secondary expression in stylar (‘female’) tissues in the outcrossing species. These observations are consistent with the outcrossing species experiencing stronger selection on female reproductive tract traits that influence male postmating performance. This study evaluates key expectations of sexual selection on postmating traits but does so in the uncommon context of flowering plants.

## Introduction

Postmating prezygotic (PMPZ) traits, which act after the delivery of male gametes and before fertilization, are critical for successful reproduction. They are also expected to be influenced by sexual selection when there is competition among genotypes for mating, with consequences for both the expression of competitive phenotypes and for the genes that underlie them (Andersson & Iwasa, 1996; Bernasconi et al., 2004). While these expectations of PMPZ traits have been examined broadly in animal systems (Firman et al., 2017; Garlovsky et al., 2023; Wigby & Chapman, 2004) they remain surprisingly poorly explored in other sexually-reproducing groups, including plants (Tonnabel et al., 2021). In flowering plants, PMPZ stages of reproduction involve direct interactions between the pollen (the ‘male’ tissue) and the pistil (the ‘female’ tissues that include the stigma on which pollen lands and germinates, and the style through which pollen tubes grow to reach the ovary). At these PMPZ stages, the success of different pollen genotypes can be influenced by both male-male pollen competition and male-female reproductive interactions (Tonnabel et al., 2021); the latter include female-mediated mechanisms that bias male gamete success, known as ‘cryptic female choice’ (Eberhard, 1996; Firman et al., 2017). Evidence supporting the action of strong selection on plant PMPZ traits comes primarily from studies of molecular evolution in genes expressed in pollen, pistils, or both tissues (Clark et al., 2006; Moyle et al., 2021). However, comparatively little attention has focused on how changing the intensity or nature of selection on PMPZ traits differentially affects the evolution of loci associated with male-male gamete competition, cryptic female choice, or both. Here our goal is to investigate PMPZ traits and loci to assess evidence for these differential effects.

One broad mechanism that could change the focus or intensity of selection on PMPZ traits is a change in mating system. Mating system shifts—primarily from outcrossing to selfing—are among the most common reproductive transitions in plants (Barrett, 2002). Because they affect the number and identity of mating partners, these shifts are predicted to change selection on PMPZ traits including those involved in pollen competition and cryptic female choice (Mazer et al., 2010). In comparison to outcrossing where pollen can be received from multiple individuals, selfing often limits male-male competition and female choice to the variation present within one individual (Cutter, 2019). Under these conditions, traits that mediate pollen competition are expected to experience relaxed selection or, if sufficiently costly, to be selected against (Cutter, 2019; Sicard & Lenhard, 2011; Wozniak & Sicard, 2018). Consistent with this, analyses of floral tissues have documented changes in gene expression (Frazee et al., 2021; Slotte et al., 2013; Z. Zhang et al., 2022) and molecular evolution (Moyle et al., 2021) accompanying shifts from outcrossing to selfing. However, because few studies investigate molecular differences in individual PMPZ reproductive tissues (although see Pease et al., 2016; İltaş et al., 2024), the association between these broad molecular patterns and changes in specific PMPZ traits and their underlying loci remains largely correlative.

Pollen tube growth rate is one such trait that mediates PMPZ interactions (Lankinen & Karlsson Green, 2015) and could be affected by mating system shifts (Mazer et al., 2010).

Because only a fraction of growing pollen tubes succeed in fertilizing available ovules within a single flower (Delph & Havens, 1998), pollen competition among different genotypes can produce strong selection for faster pollen tube growth (Delph et al., 1998; Pannell & Labouche, 2013; Snow & Spira, 1991). Moreover, factors affecting pollen tube growth rate could also be affected by male-female PMPZ sexual selection (Lankinen & Karlsson Green, 2015) where females exert genotype-specific effects on male PMPZ siring success (Fitz Gerald et al., 2014; Lu et al., 2020; Lubini et al., 2023; Marshall & Folsom, 1991; Sanchez et al., 2004). This male-male and male-female PMPZ selection on pollen growth rate is expected to be relaxed under increased selfing (Cutter, 2019; Mazer et al., 2010) so that, for example, selfing lineages have slower pollen tube growth rates compared to closely related outcrossing species (e.g., Mazer et al., 2018).

Pollen tube growth is also a trait whose molecular basis has been examined in model systems (Zheng et al., 2018). Among the contributing molecular factors, pectin methylesterases (PMEs) and their inhibitors (pectin methylesterase inhibitors (PMEIs)) are proteins with known phenotypic effects on pollen tube growth. PMEs and PMEIs influence the flexibility of pollen tube tips and shafts, modifying the degree of tissue calcification based on their relative concentration (Bosch et al., 2005; Catoire et al., 1998; Jiang et al., 2005). Moreover, since PMEs and PMEIs are expressed by both the style and the growing pollen tube (Bosch et al., 2005; Ezura et al., 2017; Pelloux et al., 2007; G. Y. Zhang et al., 2010), their influence on pollen tube growth rate depends on the combined expression of PMEs and PMEIs from both tissues. The joint ability of PMEs and PMEIs to influence pollen tube growth rate, via expression in both male and female tissues, makes them an interesting lens through which to study the contributions of male and female tissues to differential pollen performance traits between species with different mating systems.

In this study we sought to assess how loci associated with pollen tube growth (and thereby, potentially, with male-male gamete competition and cryptic female choice) are affected by a change in mating system, focusing specifically on the expression of PME and PMEI genes. To do so, we re-examined reproductive (pollen and style) and non-reproductive (leaf) gene expression data from two species in the wild tomato group—*Solanum* section *Lycopersicon*— that differ in their predominant mating system. In addition to quantifying global patterns of gene expression variation and tissue-bias, we identified reproductive-specific PME and PMEI loci and the distribution of their expression in male (pollen) and female (pistil) tissues in each species. To better interpret the relevance of detected gene expression differences for pollen performance within each species, we also quantified their pollen phenotypes *in-vivo* (growing within a female pistil/reproductive tract) and *in-vitro* (growing independent of the female pistil, in pollen growth media). Combining our observations of reproductive PME/PMEI expression with pollen tube growth rates, we develop one possible model that connects shifts in gene expression to changes in selection on male- and female-side PMPZ traits that accompany mating system changes.

## Materials and Methods

### Study species

Our analysis focused on two species in the wild tomato clade (*Solanum* section *Lycopersicum*) that vary in their mating system: *S. lycopersicum* (domesticated tomato; genotype LA3475) and *S. pennellii* (genotype LA0716). Species in this clade range from obligately outcrossing (via gametophytic self-incompatibility) through to primarily selfing (Bedinger et al., 2011). *S. lycopersicum* is genetically self-compatible and frequently self-fertilizes. *S. pennellii* is a wild tomato species that is historically genetically self-incompatible (Covey et al., 2010).

Although the specific genotype of *S. pennellii* in our experiment has recently transitioned to self-compatibility via loss of function mutations in the self-incompatibility mechanism (Covey et al., 2010), it remains facultatively outcrossing. These two species also differ in floral traits consistent with their differences in outcrossing rate (Bedinger et al., 2011; Peralta & Spooner, 2005). *S. lycopersicum* flowers are smaller, with shorter styles and inserted stigmas that promote self-fertilization; *S. pennellii* flowers are significantly larger, with a stigma and style that is exerted beyond the anthers, consistent with outcrossing (Vosters et al., 2014). A previous study of post-pollination isolating barriers between these two species (Pease et al., 2016), analyzed both pollen and stigma/style gene expression and found substantial global transcriptomic differences between them.

### RNA-seq data and gene expression quantification

Gene expression (RNA-seq) data was obtained for leaf, pollen, and stigma/style tissues from two published datasets (PRJNA245845 from Ichihashi et al., 2014, and PRJNA309342 from Pease et al., 2016). The leaf dataset (from Ichihashi et al., 2014) includes samples from four early leaf development stages (P3-P6 primordia), and two spatial positions within three of these stages (distal versus proximal samples), across three biological replicates. Because our goal was to use general patterns of gene expression in leaf tissue as a contrast for reproductive (pollen and style) tissue expression, we used the average gene expression across all seven sub-types of leaf tissue within each biological replicate. The pollen and style data were from Pease et al. (2016) which contains details of its collection. The pollen dataset included two categories, ungerminated and germinated pollen; for our analyses, we used the average gene expression across these two sub-categories, within each biological replicate. The style dataset also included two categories, unpollinated and pollinated styles; however, because pollinated styles represent a mixed tissue (both style and pollen) we only included unpollinated style data in our analysis. For simplicity, we refer to this tissue as ‘style’, although collected styles included intact stigmas so these data technically encompass both style and stigma gene expression. For both pollen and style data, we had three biological replicates.

For each of the above tissues, we used the FASTQ preprocessor fastp (Chen et al., 2018) to preprocess and quality-filter the archived raw read data. Only reads >15 bp were retained; in addition, at least 60% of bases in each read were required to have Phred quality scores greater than or equal to 25. Cleaned reads were mapped to the tomato reference genome (SL4.0) using the annotation-aware read mapper STAR (Dobin et al., 2013). Tomato genome annotation ITAG4.0 was used to create the genome index used by STAR. Read counts were estimated with featureCounts (Liao et al., 2014).

Raw read counts were normalized in two ways, depending upon the target analysis. First, analyses depending on tissue-specificity and -bias require normalization by TPM (transcripts per million) (see next section). Therefore, we applied the standard TPM normalization method, which involved determining the reads per kilobase for each gene and dividing that by the scaling factor in R. Only loci with an average TPM>=2 (across biological replicates) in at least one tissue and at least one species were included in the associated analyses. This resulted in a total dataset of 21,039 expressed loci. Second, differential expression analyses of individual loci in *DESeq2* require that normalized gene expression is quantified as the median of ratios. This was performed within *DESeq2*.

### Determining tissue-specificity and bias

We classified gene expression among tissues within species in terms of two classes of gene expression: ‘tissue-specific’ and ‘tissue-biased’ genes. ‘Tissue-specific’ genes were defined within each species dataset as those expressed solely in one tissue type (i.e., leaf, pollen, or style) and not in any other tissue (cutoffs for expression are specified below). In comparison, ‘tissue-biased’ genes were defined within each species dataset as genes whose expression was enriched in one tissue type (i.e., leaf, pollen, or style) compared to the other tissue types (as determined by the metric Tau, below). Accordingly, tissue-specific genes are a subset of tissue-biased genes. In addition, in each of our species, we also classified loci according to whether they were specific to non-reproductive or reproductive tissues (i.e., expressed in leaf tissue only, or pollen and/or style only, respectively).

Tissue-specificity and -bias were determined for each gene within each species dataset using the metric tau (Yanai et al., 2005):

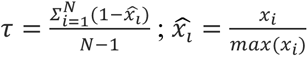

where *N* is the number of tissues and *x̂_i_* is the gene expression normalized by the maximal tissue expression of that gene. Loci with a tau value greater than or equal to 0.7 were designated as biased towards the tissue in which they were most highly expressed. All other genes were categorized as unbiased. Loci were designated as tissue-specific when they had a tau of 1.0.

### Global patterns of gene expression variation between species

To compare broad patterns of gene expression level and tissue specificity between species, we used linear models with species as a factor (Table S8). These models assessed whether there were differences between species in the average magnitude of pollen, style, or leaf tissue-specificity (i.e., the mean tau for each tissue), or in the average level of gene expression (TPM) in each of these classes of locus.

To evaluate functional classes of genes expressed within each species, genes designated as pollen, style, and leaf biased in each species were assessed for functional overrepresentation by performing a Gene Ontology (GO) term enrichment analysis in *Panther* v17 (Mi et al., 2019; Thomas et al., 2022). Each group of tissue-biased loci within each species was compared against the list of all expressed genes in our dataset. These analyses assessed enrichment in three different classifications of functional category: biological process, molecular function, and protein class. Significance was determined using Fisher’s exact tests with Bonferroni correction for multiple testing. All statistics were calculated using the PANTHER Overrepresentation Tests.

### Differential expression of individual loci between species

To identify which individual genes were differentially expressed between species, including specifically in pollen and style tissue, we used the *DESeq2* package (Love et al., 2014) in R. For each locus, normalized gene expression was quantified as the median of ratios, as required in *DESeq2*. A generalized linear model was fit to each gene in our dataset of tomato loci (N=33,562), in which expression counts were modeled as a function of species and tissue.

Replicate samples were nested within each tissue category (leaf, pollen, and style). Of all genes, N=29,999 loci were expressed at any level in any tissue. In addition, we further reduced this set to only include those loci (N=21,039) whose expression was biologically significant (i.e. >2TPM in any tissue; Table S1). For each resulting gene, two planned contrasts were performed (Table S1), specifically: pollen expression between species and style expression between species. The Benjamini-Hochberg method for multiple test correction (Benjamini & Hochberg, 1995) was applied for each contrast performed (two contrasts for each locus evaluated; post-correction p<0.05).

Among those loci significantly differentially expressed between species in pollen and style tissues, to explore functional roles of highly differentially expressed loci, we identified the 10 loci with the largest differences (TPM) between species (Table S2). Gene protein functions for these loci were evaluated based on functional annotations on the Sol Genomics Network (Fernandez-Pozo et al., 2015).

### Identifying PME and PMEI genes in the tomato genome

PME and PMEIs were identified by searching the tomato annotation ITAG4.0 for functional labels consistent with these two gene families (see Table S3 for specific terms). This search generated 76 PMEs and 32 PMEIs. For these genes specifically, we evaluated tissue-specificity and tissue-bias in each species, as well as patterns of gene expression variation among tissues and among species (Tables S4 & S5), using the same methods as applied to the larger dataset.

### Assays of pollen tube growth phenotypes

To assess rates of pollen tube growth, we performed *in-vitro* and *in-vivo* pollen germination and growth assays in five biological replicates (pollen donors) per species. We collected pollen from plants grown in the Indiana University Bloomington greenhouses, across several months (Nov 2022 to May 2023 for *in-vitro* assays; May 2023 to August 2023 for *in-vivo* assays). Pollen was collected between 8AM and 12PM via artificial vibration of floral anther cones. Pollen for *in-vivo* assays was collected fresh on the day of each assay and held in 0.2mL microcentrifuge tubes before being used. Pollen for *in-vitro* assays was collected and stored in 0.2mL microcentrifuge tubes at -20°C until use. Freezing reportedly does not affect the fertility of *Solanum* pollen, even when frozen for up to 12 months (Song & Tachibana, 2007).

Nonetheless, we also evaluated a subset of genotypes using unfrozen pollen in our *in vitro* assays, to confirm that the qualitative performance of fresh and frozen pollen did not differ in these assays. We did detect absolute differences in pollen tube growth rate between unfrozen and frozen pollen (unfrozen pollen generally grew faster; Table S6), but the relative rank of species performance in *in-vitro* assays was the same regardless of whether unfrozen and frozen pollen was used (Table S14). Before use in the *in-vitro* assays, all pollen collections from each biological replicate individual were aggregated, vortexed, and aliquoted into subsamples for use as technical replicates.

#### In-vitro assay

To assess pollen germination and growth in the absence of stylar tissue, sampled pollen was germinated on pollen growth media (PGM) of 2% sucrose, 24% PEG-4000, 0.01% borate, 20mM HEPES, 3mM calcium nitrate, 0.02% magnesium sulfate, and 0.01% potassium nitrate (Covey, Subbaiah, et al., 2010). For each assay, each pollen sample was mixed with 50 μl of PGM and placed as a droplet in one well of a 24 well cell culture plate. A sample from every biological replicate (5 per species x 2 species = 10) was included in each plate; the entire assay was repeated 5 times (5 technical replicates) in May 2023. Each replicate plate was cultured in the dark for 3 hours at 22.4°C. After 3 hours, each sample was imaged at 10x magnification (including a standard scale bar) on an AMG EVOS Fl microscope.

We used ImageJ to generate the following data from each sample image: number of viable pollen, number of germinated pollen, average diameter of pollen, and average length of pollen tubes. Pollen viability was assessed via shape; collapsed elliptic pollen grains were classified as inviable and round semi-transparent pollen grains classified as viable. In each image, pollen diameter and pollen tube growth were measured on 10 randomly sampled germinated pollen, by overlaying an 8 x 11 grid and measuring the germinated pollen that was closest to each of 10 pre-determined grid locations. Values for each image (sample) were averaged across these 10 randomly sampled pollen. Only viable pollen were included in these metrics. Technical replicates were averaged to give a single value for each biological replicate, prior to analysis.

#### In-vivo assay

To assess pollen germination and growth within stylar tissue, each of five pollen-donor individuals were assayed on pistils of three additional biological individuals (pollen recipients) from their own species. Within each species, every specific combination of pollen donor and recipient (5 donors x 3 recipients = 15 unique combinations) was repeated at least twice. Recipient plants were grown in the Indiana University Bloomington greenhouses, and assays performed between the dates May and August 2023. For each assay, unopened flowers on recipient plants were emasculated 24 hours prior to the assay, by removing the anther cone using forceps. On the following day, pollen was collected from each available (flowering) pollen donor; each emasculated style received the pollen of a single donor from their species. Pollinations were performed by coating the stigma with collected pollen held in the cap of a 0.2mL microcentrifuge tube. All pollinations were performed between 8AM and 12PM. Each pollinated pistil was collected 5-7 hours after pollination by detaching it at the point where the style meets the ovary. Collected pistils were stored in 200 μl of 25% acetic acid fixing solution at -20°C.

Aniline blue staining was used to visualize pollen tubes within each collected pistil. For each collected sample, the fixing solution was replaced with 200 μl 5M NaOH and held at 24 hours at room temperature in the dark. After 24 hours the 5M NaOH was removed, the pistil washed with 200 μl 0.1M K_3_PO_4_ and stained with 200 μl 0.01mg/mL aniline blue, 0.06M K_3_PO_4_. After staining for 5-8 hours, each pistil was imaged at 10x magnification (including a standard scale bar) under UV light on an AMG EVOS Fl microscope. Each sample required multiple images to capture the complete length of the style. Images of the same style were stitched together in Adobe Photoshop, prior to collecting phenotypic data in ImageJ. From each stitched image, we measured pollen tube length of the 5 longest pollen tubes and style length.

To analyze PMPZ phenotypes, we used linear models (T-tests and ANOVAs) in the *stats* package in R. T-tests were used to assess species differences in average germination rate, pollen tube growth rate, and pollen diameter (in *in-vitro* assays), and average style length, pollen tube growth rate, and proportion of style traveled (in *in-vivo* assays). We used follow-up ANOVAs (on pollen diameter and in vitro pollen tube growth rates) to also assess evidence for differences among individuals within each species, and (for in vivo growth rates) the effect of assay timing.

To compare the variances in *in-vitro* pollen tube growth rates between species, we used an F-test (also called a Bartlett test, or, in the *stats* package, a variance test) with species as a factor.

## Results

### Species express similar numbers of genes, but differ in the proportions of loci that are specific to pollen

Species expressed similar numbers of total genes (∼19,600; Table 1), and most (>80%) expressed genes were detected in both style and leaf tissues in both species (Table S7). For all three tissues, the fraction of genes classified as tissue-biased was also broadly equivalent between the species (Table 1), as was the proportion of style- and leaf-biased genes that were tissue-specific (i.e., loci with tau=1, Table 1). In contrast, species differed substantially in the proportion of pollen-biased genes that were pollen-specific: *S. lycopersicum* had nearly twice as many pollen-specific genes (631 of 1132 pollen-biased genes) than was observed in *S. pennellii* (340 of 1027 pollen-biased genes, Table 1, Figure 1). Consistent with this, the average magnitude of pollen bias (tau) was significantly higher in *S. lycopersicum* (linear model on all pollen-biased genes between species: p < 2e-16, Table S8).

**Figure 1:**
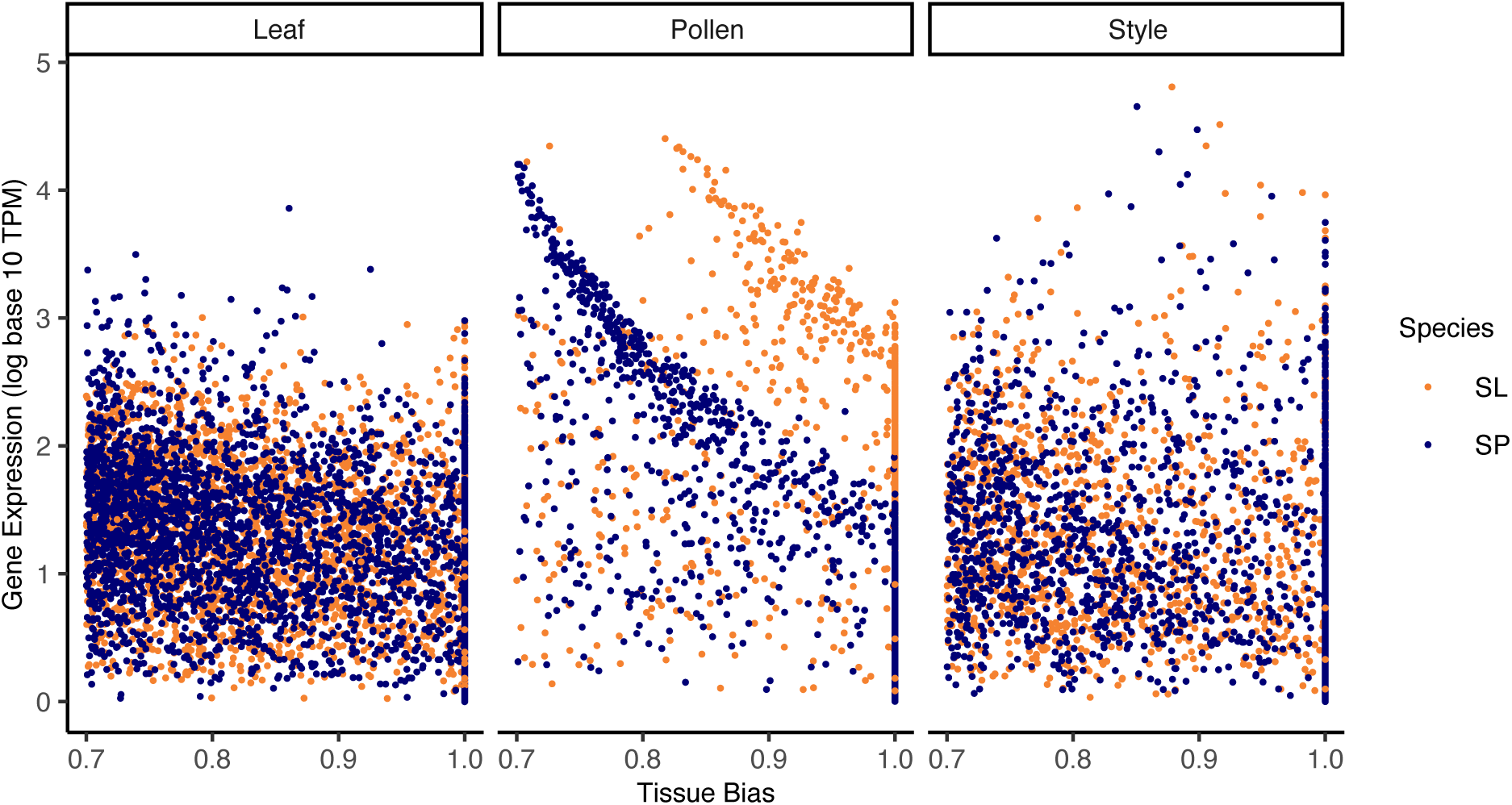
Gene expression in all leaf-biased (left), pollen-biased (middle), and style-biased (right) genes, in *S. lycopersicum* (SL, orange) & *S. pennellii* (SP, blue). Each point is an individual gene. Gene expression is normalized (in log_10_ TPM units) and tissue bias is in units of tau. See Table 1 for N in each category. The average tau of *S. lycopersicum* pollen-biased genes is significantly higher than the average tau of *S. pennellii* pollen-biased genes (linear model on tau between species: p < 2e-16, Table S8).

**Table 1:**
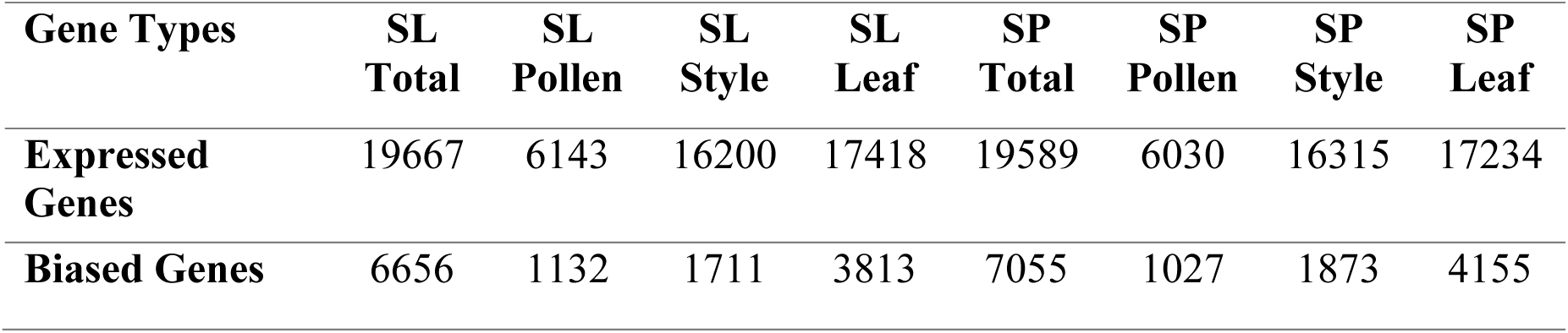
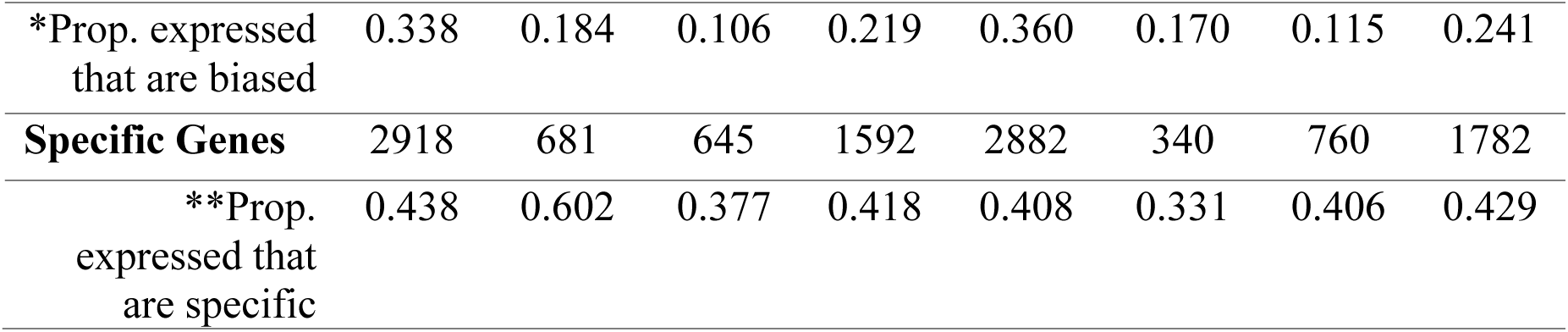
The number of genes expressed at >2 TPM in each tissue, in each species. SL = *S. lycopersicum* SP = *S. pennellii. \** within a tissue, the proportion of expressed genes whose expression is biased towards that tissue; ** within a tissue, the proportion of tissue- biased genes that were tissue-specific.

### More genes are pollen-biased or -specific in the selfing species because these loci have lower or no secondary expression in style tissue

The difference in pollen-specificity between species is not explained by differences in average pollen expression level (TPM) across all pollen-biased genes (linear model on expression between species: p = 0.754; Figure 1; Table S8). Instead, we found that genes are less pollen-biased and -specific in *S. pennellii* because these loci have significantly greater secondary expression in *S. pennellii* styles. That is, *S. pennellii* pollen-biased genes are expressed at higher levels in *S. pennellii* style tissue, compared to the stylar expression of pollen-biased genes in *S. lycopersicum* (linear model on expression between species: p < 2e-16; Figure 2; Table S8).

**Figure 2:**
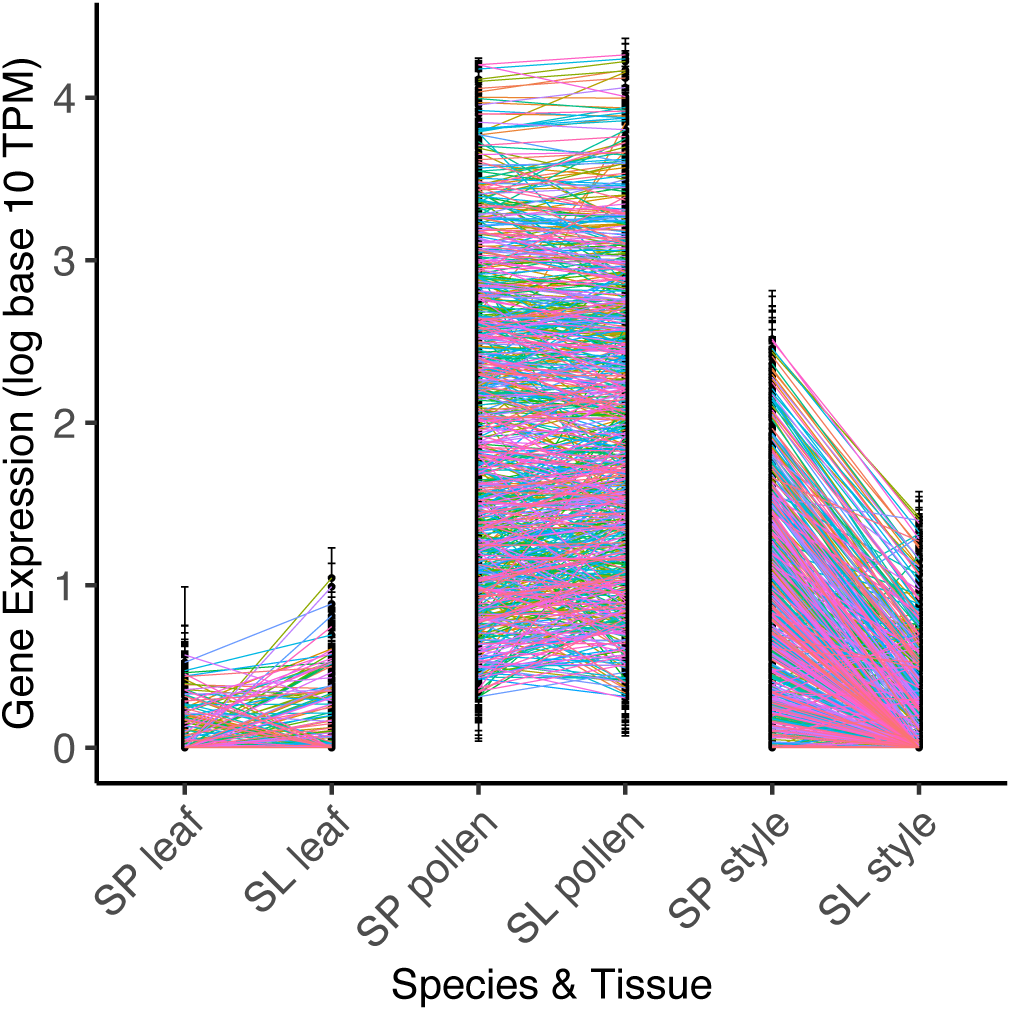
Gene expression in all pollen-biased genes with TPM >10 in pollen, by species and reproductive tissue. Gene expression is normalized (in log_10_ TPM units). Average pollen expression does not differ between species (p = 0.754, Table S8); style expression does significantly differ (p < 2e-16, Table S8). SP = *S. pennellii*. SL = *S. lycopersicum*.

Moreover, of the 482 pollen-biased genes that have secondary expression in style tissue in either species, only 11 have higher expression in *S. lycopersicum* while 471 have higher expression in *S. pennellii* (Table S9, Figure 2). Strikingly, over half of all pollen-biased genes (254 of 482) are not expressed at all in *S. lycopersicum* styles. Overall, the expression domain of pollen-biased genes is much narrower in *S. lycopersicum* compared to *S. pennellii*, a pattern only observed for pollen-biased genes.

In comparison to pollen-biased genes, *S. pennellii*’s leaf-biased genes had a significantly higher average tau than *S. lycopersicum*’s leaf-biased genes (linear model on tau between species: p = 0.003, Figure 1, Table S8). The average tau of style-biased genes did not differ significantly between species (Figure 1). This underscores that greater tissue bias is not a general feature of *S. lycopersicum* but is observed specifically in pollen-biased genes in this species.

### Pectinesterase and pectinesterase inhibitor functions are overrepresented in pollen-biased genes

Compared to all expressed genes in our dataset, our Gene Ontology analyses found several molecular functions, protein classes, and biological processes were overrepresented in pollen-, style-, and leaf-biased genes in each of *S. lycopersicum* and *S. pennellii* (Table S10). In pollen-biased genes, the pectinesterase (GO:0030599) molecular function was significantly overrepresented in both species, and the pectinesterase inhibitor (GO:0046910) molecular function was significantly overrepresented in *S. lycopersicum*. Other highly overrepresented categories for pollen-biased genes in one or both species included various actin and/or cytoskeletal protein classes (actin or actin-binding cytoskeletal protein (PC00041), cytoskeletal protein (PC00085), and non-motor actin binding protein (PC00165), as well as monovalent inorganic cation homeostasis (GO: 0006885) and multiple phospholipid and/or phosphatidylinositol molecular functions (Table S10) that are known to play an important role in pollen tissue structure (Brewbaker & Kwack, 1963) or pollen development (Lee et al., 2008) including in *S. lycopersicum* (Huang et al., 2014). The categories overrepresented in style-biased genes in both species included oxygenases (PC00177), oxidoreductases (PC00176), and metabolite interconversion enzymes (PC00262). The categories overrepresented in the leaf-biased gene group for both species were largely separate from those overrepresented in pollen- or style-biased genes, with the exception of a few general processes (Table S10).

### Reproductive genes that are highly differentially expressed between species have protein functions involving pollen signaling and development

We determined the annotated functions of the top 10 differentially expressed genes between species, in each of pollen and style tissues, for both tissue-biased and tissue-specific subsets of the data (Table S2). These included several loci with known functional roles in pollen tube growth, modification of plant cell walls, or pollen-pistil signaling. In pollen-biased genes, these included a pectin methylesterase gene (Solyc05g054360.4) which showed higher expression in *S. lycopersicum,* and a beta-D-glucosidase gene (Solyc01g009240.4) which showed higher expression in *S. pennellii.* In style-biased genes, these included a defensin gene (Solyc07g007730.4), with a GO term for peptidase inhibitor activity, that was more highly expressed in *S. lycopersicum*, and a 1-aminocyclopropane-1-carboxylate oxidase (ACC) protein gene (Solyc07g049550.3) that was more highly expressed in *S. pennellii*.

### Most reproductive-specific PMEs and PMEIs have lower style, but not pollen, expression in the selfing species

For the 76 PME and 32 PMEI genes we inferred in the tomato genome (see Methods), we determined the tissue bias and specificity of these loci (Tables S4, S5, & S11). We found 16 of 76 PMEs and 5 of 32 PMEIs were specific to reproductive tissues (i.e., expressed at >2 TPM in only style and/or pollen tissue).

Across all 16 reproductive-specific PMEs, the average level of stylar gene expression (TPM) was significantly greater in *S. pennellii* than in *S. lycopersicum* (linear model on expression between species: p = 0.0053; Figure 3; Table S8). Moreover, 15 of these 16 loci individually showed significantly higher expression (normalized median of ratios) in the style of *S. pennellii* than in the style of *S. lycopersicum* (Table S4). The remaining PME is significantly more expressed in *S. lycopersicum* style but lowly expressed in both species (5.4 vs 3.5 TPM, Table S4). Each of the five reproductive-specific PMEIs were also individually more highly expressed (normalized median of ratios) in *S. pennellii* styles than in *S. lycopersicum* styles (Table S5; Figure 3), despite their overall average expression (TPM) not significantly differing between species (linear model on expression between species: p = 0.136; Figure 3; Table S8).

**Figure 3:**
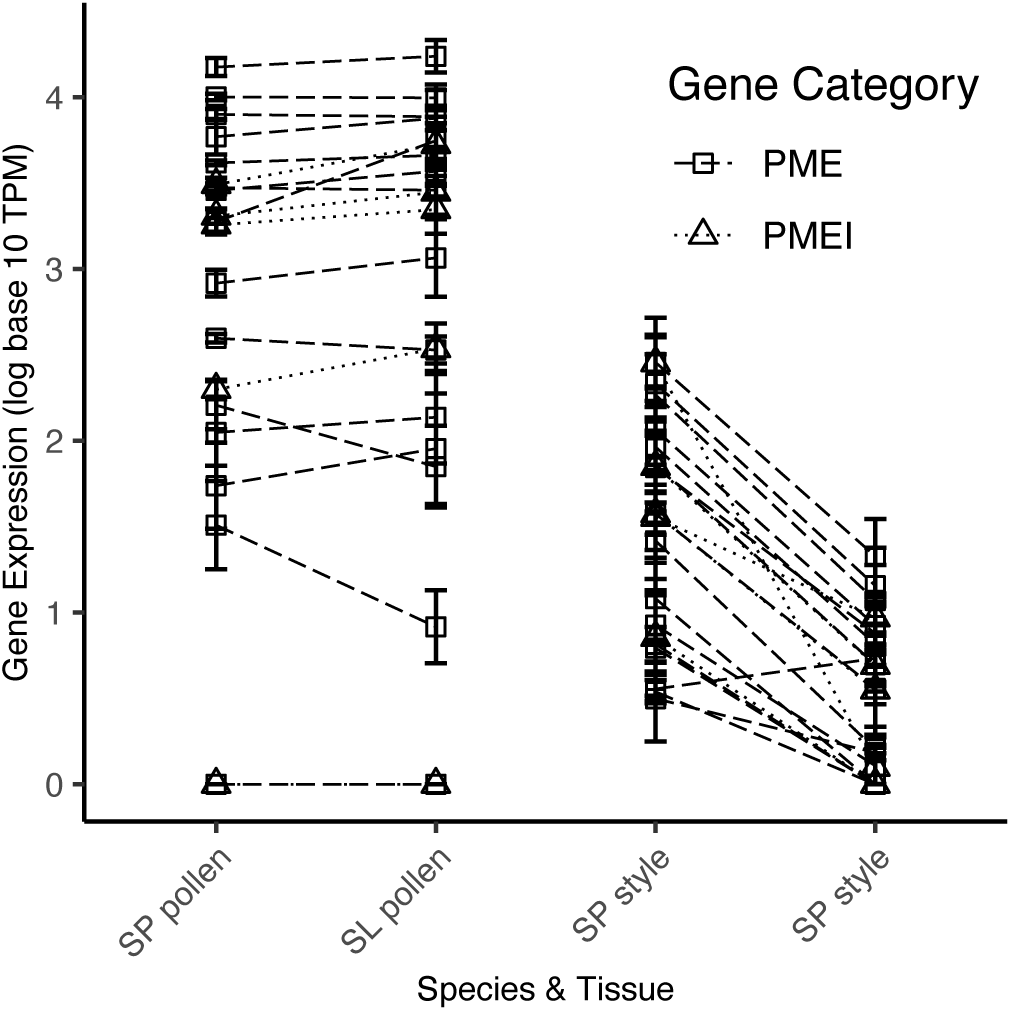
Gene expression for exclusively reproductive PMEs (N = 16) & PMEIs (N = 5), by species and tissue (Tables S4 & S5). PMEs are square, with dashed lines. PMEIs are triangles, with dotted lines. Average pollen gene expression was not significantly different between species (PMEs: p = 0.732, PMEIs: p = 0.486; Table S8). Average style gene expression was significantly different for PMEs (p = 0.0053; Table S8) but not PMEIs (p = 0.136; Table S8). Gene expression is normalized (in log_10_ TPM units). SP = *S. pennellii*. SL = *S. lycopersicum*.

In contrast to style expression, average pollen expression (TPM) across all PME or PMEI reproductive-specific genes did not differ between species (linear model on expression between species: PMEs: p = 0.732, PMEIs: p = 0.486, Figure 3; Table S8). Of the 16 reproductive-specific PMEs, one was individually more highly pollen expressed in *S. lycopersicum*, and one was more highly expressed in *S. pennellii*. No reproductive-specific PMEIs had significantly different pollen expression between species. The expression level and genomic location of all PMEs and PMEIs are given in Tables S4 and S5.

### Relative pollen tube growth rate differs between species, depending on whether pollen is growing in-vitro or in-vivo

To better interpret the role of detected gene expression differences in pollen performance traits within each species, we also quantified their pollen phenotypes in the presence (*in-vivo*) and absence (*in-vitro*) of the female style, and therefore with and without pollen-pistil interactions. Regardless of context, *S. pennellii* has larger pollen grains than *S. lycopersicum* (linear model on species, p <2e-16; Table S14), a pattern consistent among biological replicates within species (p = 0.123; Table S14).

In our *in-vitro* assays, *S. lycopersicum* pollen had significantly faster pollen tube growth rates on average than *S. pennellii* (Figure 5, linear model on species, p = 1.18e-12; Table S14). We also detected significant variation in PT growth rate among replicates within species (p = 4.05e-07; Table S14), which is explained by greater variance among biological replicates (pollen donors) in *S. lycopersicum* compared to *S. pennellii* (Bartlett test on pollen tube growth rate between species, p = 0.00013; Table S14). Unlike PT growth rates, *in vitro* pollen germination rates did not significantly differ between species (p = 0.52; Table S14).

**Figure 4:**
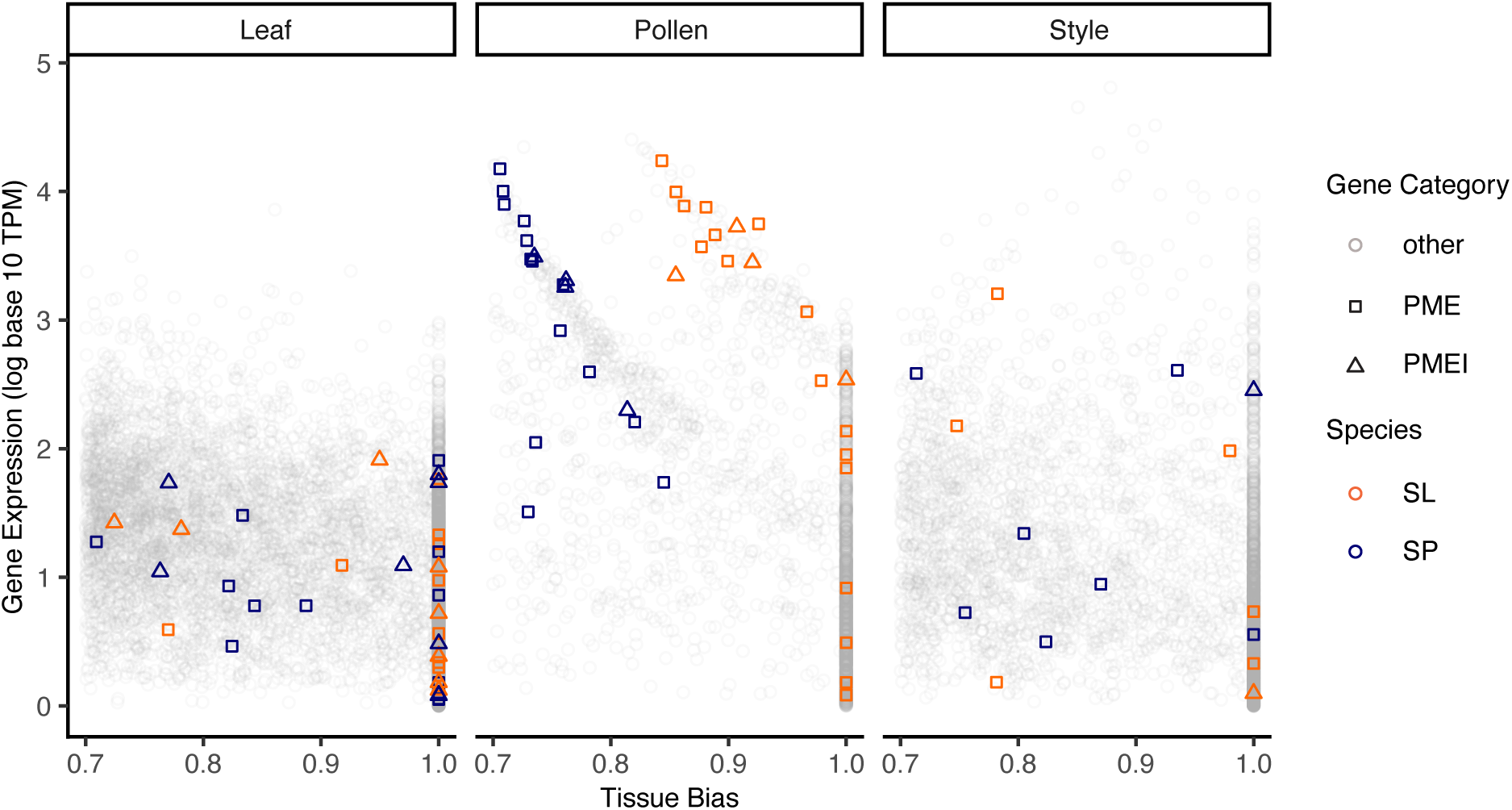
Gene expression of PMEs and PMEIs among all leaf-biased (left), pollen-biased (middle), and style-biased (right) genes, in each species. Each point is an individual gene. PMEs are squares, PMEIs are triangles, and non-PME/PMEIs are circles (and transparent gray). Gene expression is normalized (in log_10_ TPM units) and tissue bias is calculated by tau. SP (blue) = *S. pennellii*. SL (orange) = *S. lycopersicum*.

**Figure 5:**
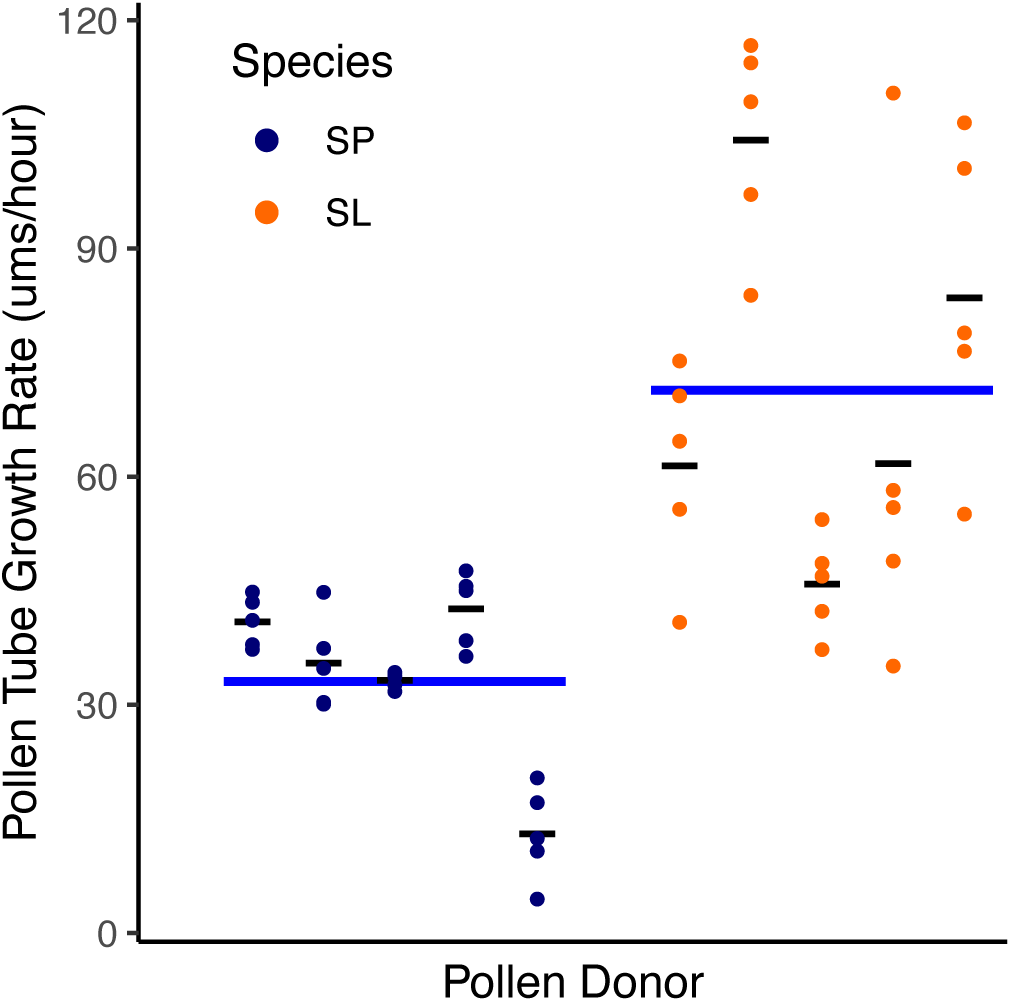
*In-vitro* pollen tube growth rate in μms/hour for *S. pennellii* (SP, blue) and *S. lycopersicum* (SL, orange). Average growth rate for each individual is shown as a thin black line. Average growth rate for each species is represented as a thick blue line. *S. lycopersicum* pollen tubes were significantly longer than *S. pennellii* pollen tubes (linear model on species, p = 1.18e- 12, Table S14c).

In contrast to *in-vitro* assays, in *in-vivo* assays we found significantly faster PT growth rates in *S. pennellii* (Figure 6, linear model on species, p = 0.0003; Table S14). We also detected significant variation in PT growth rate among biological replicates within species (linear model on biological replicates as a factor of species, p = 0.0287; Table S14) although this could not be ascribed to a specific species. Finally, because *S. pennellii* style lengths are consistently longer (linear model on species, p < 2e-16; Table S14), species did not differ in the proportion of the style traversed by pollen tubes (linear model on species, p = 0.3492; Table S14), despite *S. pennellii*’s significantly faster *in vivo* PT growth rates.

**Figure 6:**
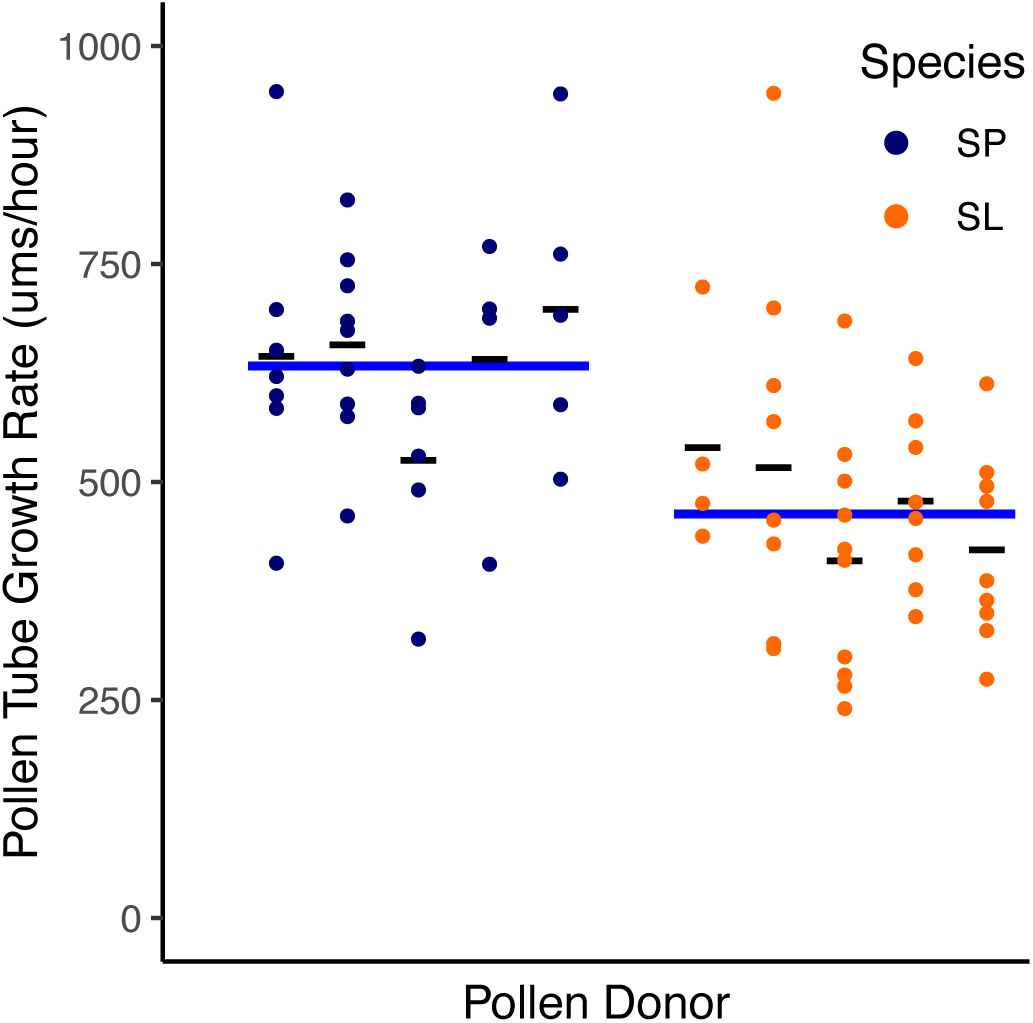
*In-vivo* pollen tube growth rate in μms/hour for *S. pennellii* (SP, blue) and *S. lycopersicum* (SL, orange), for each pollen donor. Average growth rate for each individual is represented as a thin black line. Average growth rate for each species is represented as a thick blue line. *S. pennellii* pollen tubes were significantly faster than *S. lycopersicum* pollen tubes (linear model on species, p = 0.000318, Table S14c).

## Discussion

In this study, we investigated differences in pollen, style, and leaf gene expression, particularly for PMEs and PMEIs, between historically outcrossing *S. pennellii* and selfing *S. lycopersicum*. We also observed pollen phenotypes *in-vivo* and *in-vitro* for both species, comparing pollen tube growth rates with and without the presence of the style, respectively. The most striking patterns we detected were a consistent difference between species in whether and how much pollen-biased genes were secondarily expressed in style tissue, as well as consistent species differences in pollen tube growth rates. The outcrossing species, *S. pennellii*, had greater style expression of many pollen-biased genes, including most reproductive PMEs and PMEIs. *S. pennellii* pollen also had faster *in-vivo* growth rates. In contrast, the selfing species *S. lycopersicum* had higher pollen expression of one very highly expressed PME and more rapid *in-vitro* pollen tube growth rates. Below we expand on these findings, including proposing one model for how this difference in style expression of pollen tube development genes might be due to altered selection on PMPZ traits—including via mechanisms such as cryptic female choice— between outcrossers and selfers.

### 1. Tissue-specific expression patterns identify PMPZ reproductive loci, including PME/PMEIs, that are differentially expressed between species

Our analysis identified the class of genes whose expression was biased or exclusive to two reproductive tissue types, enabling us to assess broad functional classes within each tissue, to identify PME and PMEI loci with reproductive-specific expression, and to identify differences between our focal species. Among reproductively-expressed loci (genes that are biased or exclusive to pollen and style), we found significant enrichment of several classes of pollen- performance and pollen-pistil-related functions. Among pollen-biased genes in particular, we confirmed that the pectinesterase and pectinesterase inhibitor molecular functions were significantly overrepresented in one or both species—consistent with their known role in pollen tube growth and development. Other overrepresented categories in one or both species are broadly consistent with essential cellular functions in our reproductive tissues (see Results).

We also identified the subset of *Solanum* PMEs and PMEIs (16 and 6, respectively) that are reproductive specific (Tables S4 & S5)—and could thereby have potential functions in pollen tube growth modulation. Almost all of these loci also showed significant stylar expression differences between species, suggesting their roles in the style have diverged between *S. pennellii* and *S. lycopersicum*. In contrast, only two PMEs differed in pollen expression between species, although one of these—a pectin methylesterase gene (Solyc05g054360.4) with elevated expression in *S. lycopersicum*—was among the top 10 most differentially expressed pollen-biased genes between species (Table S2).

Beyond PMEs and PMEIs, species were also significantly differentiated for loci with other known pollen performance and pollen-pistil functions (Table S2). Highly differentially expressed style-specific genes included a 1-aminocyclopropane-1-carboxylate oxidase (ACC) protein gene (Solyc07g049550.3). ACC proteins have both previously been associated with pollen tube directional signaling in *Arabidopsis thaliana* and other systems (Higashiyama, 2010; Mou et al., 2020), as well as stimulating pollen tube growth in *S. lycopersicum* (Althiab-Almasaud et al., 2021).

Together, these results indicate an enriched expression of known and putative pollen- performance and pollen-pistil loci—including multiple PMEs and PMEIs—in the reproductive tissues of these two species, as well as evidence that numerous of these PMPZ loci have significantly diverged between species in their expression.

### 2. Differences in pollen-biased gene expression between male (pollen) and female (stylar) tissues are consistent with altered selection on pollen-pistil interactions by mating system

Among all the tissue- and species-specific patterns we observed, the most general and consistent difference was a narrowed expression domain for pollen-biased genes in *S. lycopersicum* — the selfing species—compared to a broader expression domain in the outcrossing lineage. Over 95% of pollen-biased genes with any secondary expression in style tissue were more highly expressed in *S. pennellii* styles compared to *S. lycopersicum* styles (Table S9, Figure 2). Of these, nearly 300 loci are not expressed in *S. lycopersicum* styles at all. This pattern was recapitulated in reproductive PMEs and PMEIs specifically, where most loci were pollen-biased and had lower style, rather than pollen, expression in the selfing species (Tables S4 and S5).

Observing large differences in floral gene expression between species with different mating systems is not unprecedented, having been previously reported in *Arabidopsis*, *Capsella*, and *Collinsia* (Frazee et al., 2021; Slotte et al., 2013; Z. Zhang et al., 2022), as well as between these specific *Solanum* species (Pease et al., 2016). However, of the floral expression changes attributed to stylar tissue, many affect style-biased or -exclusive functions—not, as we observe here, the stylar expression of pollen-biased genes. For instance, the transition from self-incompatibility to self-compatibility involves the mutational loss of specific stylar proteins that govern the self-incompatibility reaction (Bedinger et al., 2017; Kondo et al., 2002; Stone, 2002) as has previously been reported in *S. lycopersicum* (e.g., Bedinger et al., 2011; Pease et al., 2016)) and other self-compatible species (e.g., Slotte et al., 2013). Style gene expression changes could also arise from floral development changes that often accompany increased selfing— especially reduced style length. The significantly shorter style of *S. lycopersicum* (Vosters et al., 2014), and the reduced or abolished function of specific stylar growth proteins (Pease et al., 2016), have also previously been reported in this species. Unlike these effects, mating system differences in style gene expression between species have not previously been connected specifically to pollen-related functions, although İltaş et al., (2024) recently reported differences in 25 pollen-expressed loci within pollinated pistil tissue when comparing two *Arabidopsis lyrata* populations with different mating systems.

What factors associated with mating system might specifically alter the dynamics acting on style expression of pollen-biased genes, including PMEs and PMEIs that modulate pollen tube growth? One such factor is the nature and intensity of selection acting on pollen performance traits (Cutter, 2019; Mazer et al., 2010, and see Introduction). In particular, by reducing the number and diversity of pollen donors, transitions from outcrossing to selfing are expected to decrease the benefit of stylar mechanisms that amplify discrimination among alternative pollen donors (cryptic female choice). This altered selection on stylar-mediated pollen performance would be observed as reduced style-specific influence on pollen tube growth in lineages with higher selfing. This is the pattern we observe in our analysis: lower stylar expression of pollen-biased genes in *S. lycopersicum* is consistent with a reduced allocation of stylar proteins to pollen tube development and growth, compared to *S. pennellii*. Conversely, higher expression of pollen-biased loci—including PMEs and PMEIs—in the style of *S. pennellii* indicates a greater stylar contribution to modulating pollen tube growth in this outcrossing species. This is similar to changes in gene expression observed in pollinated pistils between *Arabidopsis lyrata* populations (İltaş et al., 2024), a difference also attributed to differential selection for cryptic female choice. Resource reallocation is commonly observed following transitions to selfing, including shifts in investment from pollen to ovule production (as evidenced by reduced pollen-ovule ratios in more selfing species; Cruden, 1977; Mione & Anderson, 1992; Wozniak & Sicard, 2018). Our proposal here hypothesizes a similar realignment, whereby sex allocation to stylar mechanisms of pollen tube growth is reduced in the selfing species.

### 3. In vivo pollen performance differences are consistent with mating system effects on PMPZ pollen-pistil traits

If, as we infer, expression differences between species reflect differences in stylar contributions to pollen tube growth, these species should exhibit differential pollen performance in the presence of their own style. Consistent with this, in our *in-vivo* assays we observed significantly faster pollen tube growth rates in *S. pennellii* (Figure 6), whose stylar expression of pollen tube development genes was significantly higher, compared to *S. lycopersicum* (Figure 2). Moreover, we know this difference in pollen performance is not intrinsic to pollen (or factors like larger pollen size; e.g. McCallum & Chang, 2016) alone, because the direction of differential pollen performance is reversed in the absence of stylar contributions to pollen tube development—when pollen is cultured *in-vitro*. Under these conditions, *S. lycopersicum* pollen grows comparatively faster (Figure 5). The specific molecular details of this differential stylar influence over pollen performance remain to be clarified. However because the interaction of PMEs and PMEIs is known to control the equilibrium of pectin methylesterification in growing pollen tubes (Bosch & Hepler, 2005; and see Introduction) one possibility is that the higher joint expression (across both pollen and style) of PMEs, PMEIs, and possibly other pollen growth factors enables more rapid *S. pennellii* pollen tube elongation within *S. pennellii* stylar tissue, compared to *S. lycopersicum* pollen growing in *S. lycopersicum* styles.

Under this model, our observation that *S. lycopersicum* has comparatively faster growth rates in the absence of a style (*in-vitro*, Figure 5) conversely suggests that its pollen might have higher intrinsic production of pollen growth factors compared to *S. pennellii*. We did not generally observe elevated pollen-biased gene expression in *S. lycopersicum* pollen, but detected one notable exception: among the top 10 most differentially expressed pollen-biased genes (Table S2) is one PME (Solyc05g054360.4) with 3-fold higher expression in *S. lycopersicum* pollen (average TPM of 5596 vs. 1186 in *S. pennellii*; Table S15). The biological significance of this locus for differences in species intrinsic *in-vitro* pollen tube performance remains to be determined; however, it is intriguing to speculate that it might represent a compensatory response to the lower expression of PME genes in the *S. lycopersicum* style. Ultimately, polarizing the timing and direction of evolution of these specific expression differences between species—as well as those observed in stylar tissue—would benefit from additional gene expression data from other closely related species that also vary in their predominant mating system.

### 4. Implications for postmating prezygotic trait evolution under different mating systems

Changing mating strategies can relax old constraints, and potentially impose new ones, on many traits that influence the number and identity of reproductive partners (Barrett, 2002; Cutter, 2019; Mazer et al., 2010). Our two focal species differ in reproductive traits that are known to be associated with mating system transitions, including changes in floral size, flower allometry (stigma exsertion), and loss of specific stylar pollen-rejection mechanisms (Bedinger et al., 2017; Chalivendra et al., 2013; Covey et al., 2010; Jewell et al., 2020; Vosters et al., 2014).

Our analysis here indicates that altered stylar expression of pollen-associated loci, with associated changes in the rate and stylar-independence of pollen tube growth, are also among the significant reproductive changes that differentiate these two species.

Our inference is that these differences are due to changes in the kind or strength of selection acting on postmating prezygotic traits—specifically style-mediated elements of pollen-pistil interactions, as similarly inferred in İltaş et al., (2024). The precise eco-evolutionary conditions responsible for this observed difference remain to be definitively established. As proposed above, reduced sex allocation to stylar mechanisms of pollen tube growth could result from relaxed selection on cryptic female choice in more selfing lineages. If so, this implies that stylar-expressed pollen growth factors are normally involved in exerting this choice in outcrossing species. PME has been implicated in mate choice in the form of PMPZ reproductive isolation between maize species (Wang et al., 2022) so it is known to mediate stylar mechanisms of discrimination among pollen of different genotypes. Higher style expression of pollen growth genes in *S. pennellii* may enable greater fine-tuning of *in-vivo* pollen performance, for instance by amplifying the growth of pollen whose intrinsic expression of PMEs and PMEIs is best matched to intrinsic stylar expression of these same factors. Nonetheless, these style-mediated pollen-pistil interactions might also be shaped by several other dynamics (e.g. Harder et al., 2016) that could differ between outcrossing versus selfing species. For instance, stylar promotion of pollen tube growth might better ensure the ovary receives sufficient pollen tubes to fertilize ovules, under conditions of strong pollen limitation; conversely, pollen limitation might be alleviated by increased selfing, reducing the need for stylar supplementation of pollen growth (Harder & Aizen, 2010). Note that these style-mediated mechanisms need not be mutually exclusive. Promoting (compatible) pollen growth under conditions of pollen limitation and exercising cryptic female choice among pollen might be achieved by the same or similar mechanisms of stylar inputs into pollen growth.

Regardless of the specific reproductive conditions responsible, our analysis indicates that their consequences have primarily played out as changes in the gene expression profiles of styles. While other studies have also noted changes in pollen-related gene expression between mating systems using whole flower transcriptomes (Frazee et al., 2021; Slotte et al., 2013; Z. Zhang et al., 2022), or pollinated pistil gene expression (İltaş et al., 2024) these did not localize these changes to the secondary expression of pollen-biased genes specifically in styles. In comparison to our detected style expression differences, evidence of systematically altered pollen gene expression is less clear in our study, although elevated expression of select pollen growth factors in *S. lycopersicum* might itself be a response to reduced stylar expression of these genes. This largely ‘female’ response is particularly interesting, as one of its major phenotypic consequences appears to be substantial differences in male reproductive performance in the form of pollen tube growth rates. Pollen tube growth rates have direct consequences for siring success when pollen of different genotypes are competing, so rate variation is often attributed to differences in the strength of selection on pollen competition (Mazer et al., 2010; Snow & Spira, 1991, 1996), including between closely related species that differ in mating system (e.g., Mazer et al., 2018). Our analysis here suggests that our species differences in *in-vivo* pollen performance are instead primarily the consequence of shifted stylar (‘female’) investment in pollen (‘male’) performance, a pattern that would be hidden in the absence of information on style and pollen gene expression variation. Our inference underscores that a complete understanding of evolutionary variation in PMPZ traits, including ‘male’ traits like pollen tube growth, requires the joint consideration of both male and female postmating reproductive dynamics, including how each might be shaped by changes like mating system transitions.

## Conclusions

Mating system differences between two Solanum species are associated with widespread changes in pistil gene expression, particularly of pollen-biased loci including pollen tube development and growth genes. These differences appear to have attendant changes in pollen tube growth rates in the presence and absence of stylar factors. We infer that these species differ in style-specific control of pollen performance *in-vivo* as a result of reduced sex allocation to stylar mechanisms of pollen tube growth in the selfing species, consistent with relaxed selection on cryptic female choice.

## Data Availability

Raw transcriptome data were obtained from NCBI SRA BioProject# PRJNA309342. Data (including supplemental tables, read counts, *in-vitro* & *in-vivo* images, and processed data used in R analyses), shell scripts, and R markdowns used for analyses are available on GitHub (https://github.com/biewerheisler/SolanumPenLycGeneExpressionAnalysis).

## Acknowledgments

Thanks to the staff of the IU Biology Greenhouse facility for their assistance with plant growth and maintenance.

